# Using absorbance detection for hs-SV-AUC characterization of adeno-association virus

**DOI:** 10.1101/2024.03.27.587092

**Authors:** Nicholas R. Larson, George M. Bou-Assaf, Thomas M. Laue, Steven A. Berkowitz

## Abstract

Data are presented demonstrating that absorbance detection can be used during high-speed sedimentation velocity analytical ultracentrifugation (hs-SV-AUC) experiments to accurately characterize adeno-associated virus (AAV) drug products. Advantages and limitations of being able to use this detector in this specific type of SV-AUC experiment are discussed.

## Technical Note

Compared to lower-speed protocols, the high-speed sedimentation velocity analytical ultracentrifugation (hs-SV-AUC) protocol provides improved resolution, shorter run times, and higher sample throughput for characterizing large, complex supra-macromolecular assemblies [1]. A key feature of hs-SV-AUC is the need to use interference (IF) detection to achieve rapid data acquisition and avoid the adverse effects from steep, fast-moving sample boundaries. Here we investigate the accuracy of the hs-SV-AUC protocol when using absorbance detection. The motivation for this undertaking was the desire to use spectral data to: 1) characterize the compositional heterogeneity of AAV-based therapeutics [2-4], to better understand their compositional variability and 2) improve their batch-to-batch manufacturing consistency.

The hs-SV-AUC protocol [1] was modified slightly to allow acquisition of both IF and 230 nm absorbance data. In particular, the run temperature was decreased from 10 °C to 5 °C, thus allowing more scans/cell and reducing the chance for convection [5]. The protocol was performed on four different days using fresh aliquots from the same AAV sample. Data from both detectors for each run were processed using SEDFIT [6] (version 15.01b) to generate sedimentation coefficient distribution plots, c(*s*) vs *s*, where, *s*, is the sedimentation coefficient in Svedbergs, and c(*s*) is the concentration of material having a particular *s*. This model assumes the AAV samples consist of a distribution of discrete, ideal components. The distributions from each detector were compared qualitatively and quantitatively to assess differences. Since the concentration response factor for proteins and nucleic acids for absorbance detection at 230 nm an IF (refractometric) detection is approximately the same [3,7,8], differences in the protein/nucleic acid ratio for different AAV components will result in consistent signal ratios for the two detectors. Hence, if accurate absorbance data is acquired using this hs-SV-AUC protocol, nearly identical area-normalized c(*s*) vs *s* plots will be obtained (qualitative comparison). In addition, fractional amount of the various key regions of the distribution, corresponding to specific components (low molecular weight (LMW), empty, partially-filled, full and high molecular weight (HMW) AAV material), also should be the same for both detectors (quantitative comparison). Note, the HMW region may include monomeric AAV particles that encapsulate more nucleic acid material than the gene of interest (or payload).

Table 1 summarizes the fractional amount for each component determined from each detector. These results reveal good agreement between the IF and absorbance detectors, even for species present at low levels (e.g., LMW), which are difficult to quantitate [9-11]. Moreover, when the four c(*s*) vs *s* plots for each detector were averaged, area-normalized and then superimposed, the distributions are in good qualitative agreement (Figure 1A). Nevertheless, there is a slight overall offset in *s* values in the 230 nm plot relative to the IF plot. This small shift, best seen in the case of the full AAV peak, appears roughly as a 0.8 S or 1.3% higher *s* value in the 230 nm distribution compared to the IF distribution. In fact, if one decreases all the 230 nm S values in Figure 1A by 1.3%, a noticeably better overall alignment with the resulting IF distribution is obtained, Figure 1B. The same offset is observed in all four AAV hs-SV-AUC runs using the same Optima (Supplemental Table S1). Furthermore, this same shift is found for a second Biogen Optima and has been observed on other Optima instruments (unpublished information), suggesting there is a common origin for the problem. Two possibilities are: 1) differences in radial calibration [12-14] or 2) differences in how ω^2^t is calculated between these two detectors. At present, we do not know which detector provides more accurate S values, but Optima users should be aware of and test for this possibility. Despite the nuisance of this offset, the quality of agreement between 230 nm and IF data supports the conclusion that absorbance detection may be used for AAV hs-SV-AUC.

**Table 1.** ^a^Comparison of the key attributes of the same AAV sample from four hs-SV-AUC experiments where both absorbance, 230 nm, and Interference, IF, detection was used.

| **Attribute** | **<sup>f</sup><IF> (Stdev)** | **<sup>f</sup><230 nm> (Stdev)** | **Average** | **Stdev** | **<sup>g</sup>CV** |
| --- | --- | --- | --- | --- | --- |
| **<sup>b</sup>%LMW** | 0.73 (0.46) | 0.66 (0.30) | 0.70 | 0.05 | 6.6% |
| **%Empty Peak** | 7.0 (1.7) | 6.9 (1.5) | 6.9 | 0.1 | 1.8% |
| **<sup>c</sup><sub>S<sub>5</sub></sub>, S** | 41.3 (0.4) | 42.3 (0.1) | 41.8 | 0.7 | 1.7% |
| **<sup>d</sup>%Partials** | 14.2 (1.2) | 13.4 (1.2) | 13.8 | 0.6 | 4.3% |
| **%Full Peak** | 67.5 (4.2) | 67.9 (4.0) | 67.7 | 0.3 | 0.5% |
| **<sub>S<sub>5</sub></sub>, S** | 62.7 (0.2) | 63.4 (0.1) | 63.0 | 0.5 | 0.8% |
| **<sup>e</sup>%HMW** | 10.5 (4.9) | 11.1 (4.2) | 10.8 | 0.4 | 3.9% |

<sup>a</sup>Experiments were conducted on the same Optima analytical ultracentrifuge equipped with absorbance and interference detection at 45K rpm, at 5 °C at about the same AAV concentration (~1.0 OD<sub>230nm</sub> corresponding to ~0.25 fringes).
<sup>b</sup>All AAV material moving slower than the AAV empty peak
<sup>c</sup>Sedimentation coefficient at 5 °C
<sup>d</sup>All AAV material moving between the AAV empty and full peak
<sup>e</sup>All AAV material moving faster than the full AAV peak
<sup>f</sup>Average value from the four hs-SV-AUC experiments
<sup>g</sup>Coefficient of Variance

**Figure 1.**
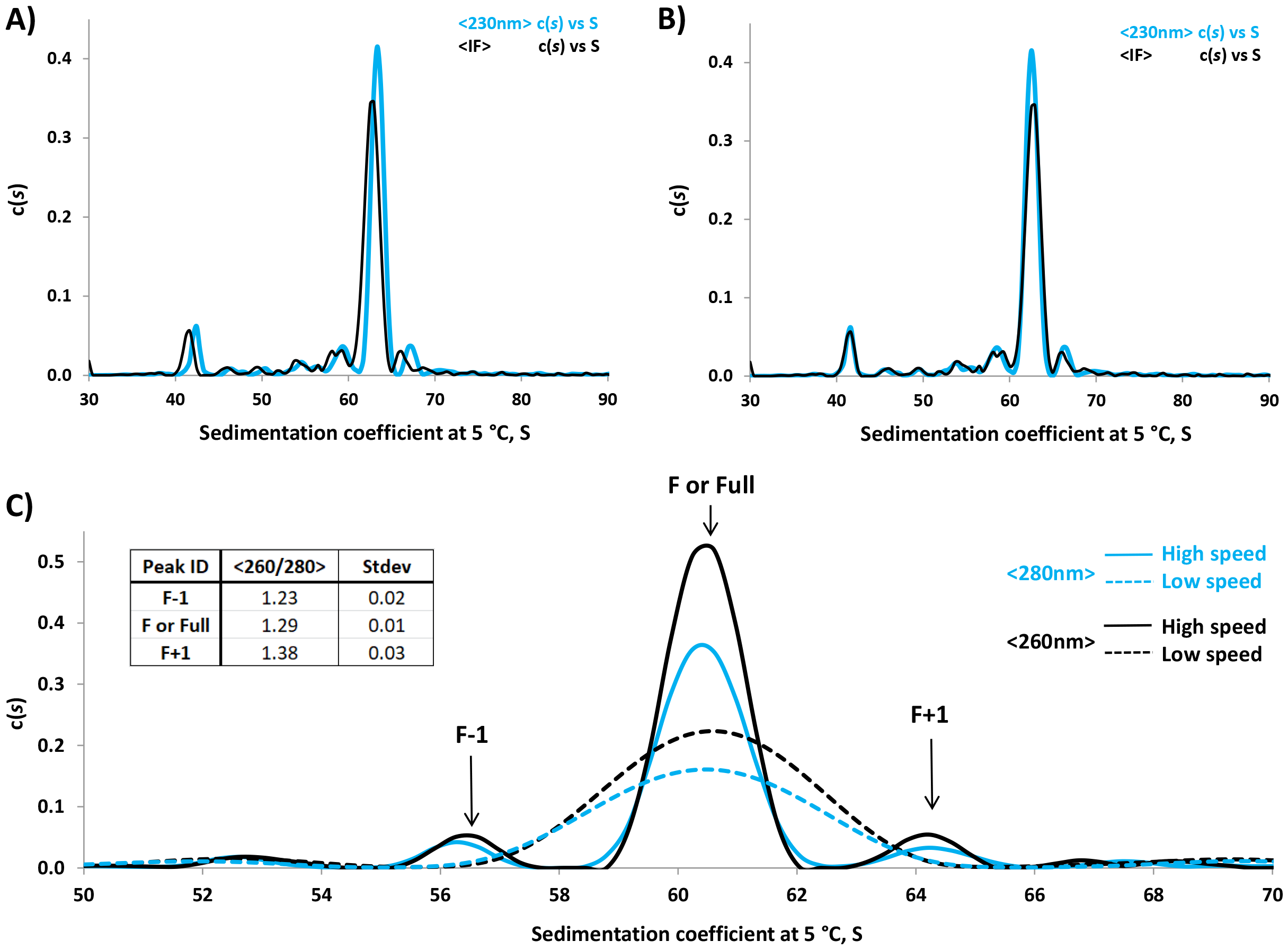
Qualitative comparison of the absorbance (230 nm) and interference (IF) area-normalized AAV hs-SV-AUC c(*s*) vs *s* plots (parts A & B) generated from hs-SV-AUC experiments with dual detection and comparison high speed vs low speed c(*s*) vs *s* plots generated from experiments where only absorbance data was acquired at 260 and 280 nm for the same AAV sample (part C): **A)** the overlay of the average absorbance (230 nm) c(*s*) vs *s* plot (light blue) with the average interference c(*s*) vs *s* plot (black) generated from four separate hs-SV-AUC experiments (noting that 230 nm and IF data was collected from the same single cell at 45K rpm and 5 °C, where absorbance data scans were collected every 20 sec and interference data was collected every 5 sec, while data from both detectors were analyzed over the same radial-time window, of 6.4-7.0 cm and ∼1100 sec, using S_min_ and S_max_ values of 30S and 130S, with all other analysis parameters set as indicated in reference [1]), **B)** same overlay plots shown in part “**A”**, but the absorbance’s x-axis S values were reduced by 1.3% to bring the S value for the absorbance full AAV peak into alignment with the S value for the interference full AAV peak and **C)** the overlay of the average absorbance 260 nm, black, and 280 nm, light blue, c(*s*) vs *s* plots localized to the area containing the AAV full peak (F) for the low speed SV-AUC experiments (dashed lines, 15K rpm) with the corresponding hs-SV-AUC experiments (solid lines, 45K rpm) at 5 °C, showing the unique presences of peaks F-1 and F**+**1 in the hs-SV-AUC data with different 260/280 peak area ratio values from the AAV full peak (F) as indicated in the table inset. Low speed data scans were collected every 75 sec (resulting in data scan spacing for each wavelength of 150 sec) and analyzed over the radial-time window of 6.2-7.1 cm and ∼10,500 sec, while hs-SV-AUC data was collected every 18 sec resulting in data scan spacing for each wavelength of 36 sec) and analyzed over the radial-time window of 6.2 -7.1 cm and ∼1280 sec. S_min_ and S_max_ values for both rotor speeds were set at 10S and 100S respectively. All other analysis parameters were set as indicated in reference [1] for both rotor speeds. Note, due to the use of a different Optima instrument in parts A & B than that used in part C, a small difference in the sedimentation coefficient values of about a 4% was seen between parts A & B and part C. A bulk of this difference can be attributed to observed temperature differences, of several tenth of a °C between these two Optima instruments. No attempt was made to account for this temperature difference between the two Optima units.

Using absorbance-based hs-SV-AUC at 260 and 280 nm, the compositional differences (protein to nucleic acid ratio) of AAV components were explored. The same AAV sample and protocol employed in Figure 1A was used in three separate runs, each conducted on a different day, where *only* absorbance data at 260 and 280 nm were collected. However, this work was conducted on a different Optima instrument than the one used in the interference/absorbance (230 nm) experiment above. Using a single cell, we were able to collect ∼30 scans at each wavelengths, enough to provide reliable c(*s*) vs *s* plots [15, 16]. For comparison, three low speed (15K rpm) SV-AUC experiments also were conducted on different days using the same second Optima instrument, AAV sample, wavelengths, and temperature. A comparison of the c(*s*) distribution attributes, listed in Table 1, for the two speeds are shown in Supplemental Table S2. An overlay of the average 260 nm and 280 nm c(*s*) vs *s* distributions for the two speeds is shown in Figure 1C.

As observed previously [1], the hs-SV-AUC protocol improves the c(*s*) resolution significantly over low-speed data (Figure 1C). Furthermore, the higher-resolution provides new, important information about the sample. In particular, hs-SV-AUC reveals that what originally was considered to be the full AAV peak at low speed (dashed traces in Figure 1C for 260, black, and 280 nm, light blue), appears to split into three distinct peaks, F-1, F or Full, and F+1, at high speed (solid traces in Figure 1C for 260, black, and 280 nm, light blue). These three peaks are observed consistently for both absorbance and interference data, lending validity to their uniqueness. Nevertheless, questions still lingered as to whether peaks F-1 and F+1 are fitting artifacts or truly unique AAV solution components. However, the A_260_/A_280_ peak area ratios, (Table insert, Figure 1C) provide a second test of the uniqueness of peaks F-1, F or Full and F+1. As shown in this table, the average A_260_/A_280_ peak area ratios increase from 1.23 ± 0.02 for peak F-1, to 1.29 ± 0.01 for peak F or Full and 1.38 ± 0.03 for peak F+1. A two-sample t-test shows these ratios are statistically different at a > 95% confidence level. Thus, the chemical composition information is orthogonal evidence that peaks F-1, F or Full and F+1 are unique AAV solution components. What was thought to be the homogeneous “full AAV” particles comprising 82 ± 1% of the sample (the average of 260 nm and 280 nm data, Table S2) by low-speed SV-AUC, is in fact only 75 ± 1% of the sample at high speed (Table S2). In other words, using the hs-SV-AUC protocol prevents the overestimation of the active pharmaceutical ingredient (API), full AAV particles, and the underestimation of product-related impurities in the final drug product. Consequently, the ability to collect data at more than one wavelength at high speed provides more confidence and potential identification of the different components present in an AAV sample, allowing for enhanced product characterization compared to low-speed analysis.

An additional positive feature of using absorbance detection when doing AAV hs-SV-AUC is the expanded accessible *s* range. Due to the prolonged calibration time needed for the Optima IF detector, the accessible *s*_*10*_ range was limited to ∼25 to ∼150 S [1]. However, using only absorbance detection, the upper *s*_*10*_ limit for the Optima can exceed 300 S (which is made even larger when running at 5 °C due to the slower migration resulting from the increased solution viscosity [1]), while in the case of the lower *s*_*10*_ limit, values much lower than 25 S can easily be achieved depending on one’s willingness to extend data collection time.

In summary, absorbance detection can be used for the analysis and characterization of AAV products when doing hs-SV-AUC with the Optima analytical ultracentrifuge. However, at 45K rpm, AAV samples sediment quickly (within 20 min) and the collection of sufficient number of absorbance scans becomes challenging. In the case of dual detection using absorbance and IF, the associated challenges are even more daunting, as noted previously [1]. Nevertheless, if one reduces the number of scans/cell from ∼50 to ∼20-30 [15,16], operates at 5 °C, and uses only absorbance detection, sufficient data from two, or possibly three, cells may be acquired. Using intensity detection data from four, or possibly six, samples may be acquired [17]. At the moment, for dual-wavelength detection, we recommend collecting data on only one cell. Clearly, despite the throughput limitations, the added ability to acquire spectral information with hs-SV-AUC is valuable.

