## Supplementary figures and images for "Using absorbance detection for hs-SV-AUC characterization of adeno-association virus"

### Supplemental Material

**Supplemental Material**

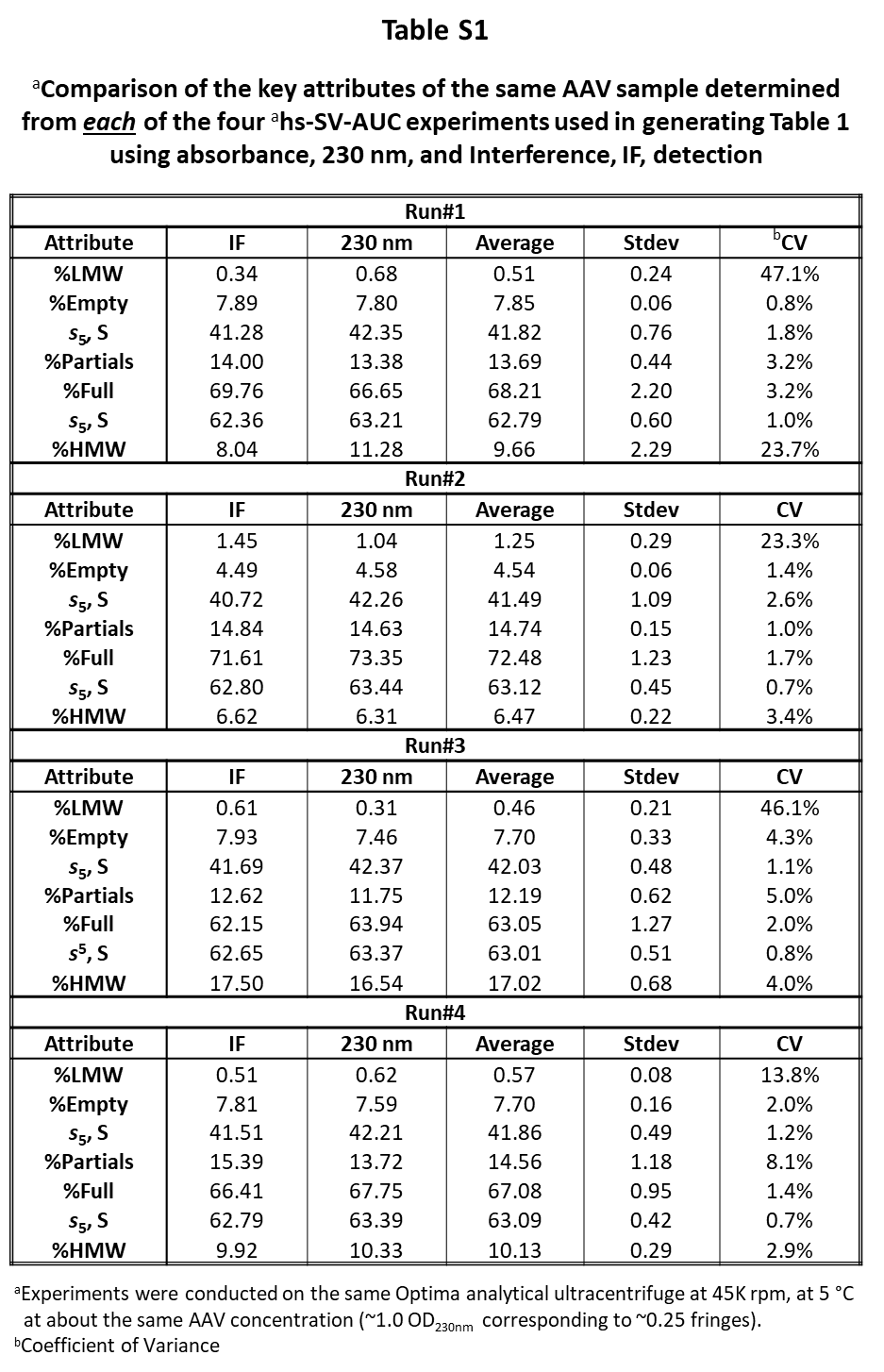

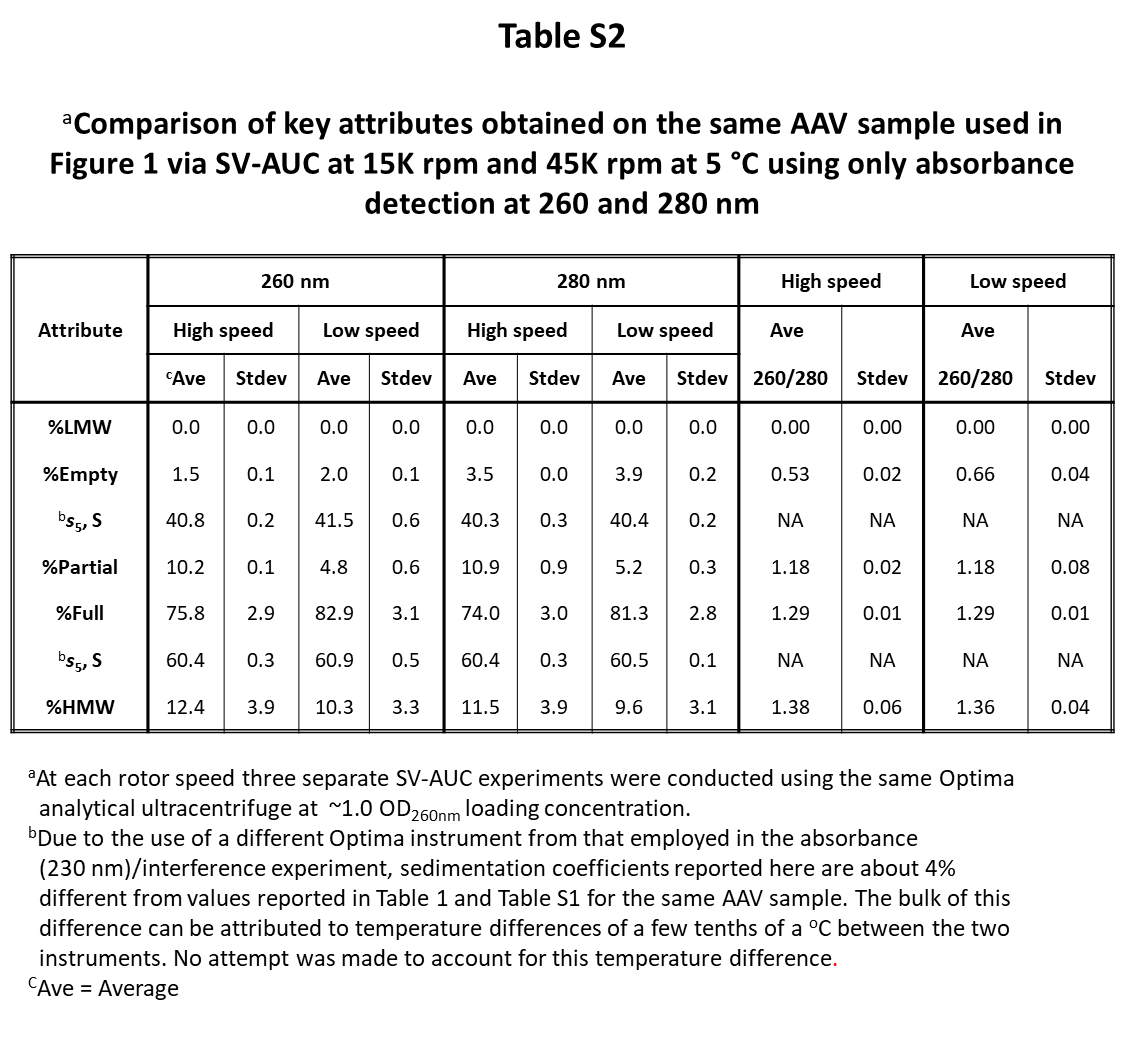
